# Relation-Aware Graph–Hypergraph Networks for Antibody Sequence–Structure Co-Design

**DOI:** 10.64898/2026.10.12.758263

**Authors:** Jinzhu Wei, Shuijun Zhang, Hongliang Duan, Leyi Wei, Jingjing Guo

**Affiliations:** Centre in Artificial Intelligence Driven Drug Discovery, Faculty of Applied Sciences, Macao Polytechnic University, Macao, 999078, China; College of Life Sciences, Nanjing Agricultural University, Nanjing, 210095, China; Engineering Research Centre of Applied Technology on Machine Translation and Artificial Intelligence, Macao Polytechnic University, Macao, 999078, China

**Author notes:** Corresponding author (J. Wei); (S. Zhang); (H. Duan); (L. Wei); (J. Guo).

**Keywords:** graph-hypergraph networks, *SE*(3)-equivariance, multi-body feature modeling, sequence-structure co-design, antibody design

## Abstract

The design of antibody complementarity-determining regions (CDRs) is pivotal for therapeutic development, yet remains challenged by the complexity of the antibody–antigen interface. Existing methods, which largely rely on pairwise relationships, struggle to capture intrinsic multibody interactions and global structural constraints. To address this limitation, we propose the *Relation-Aware Graph–Hypergraph Network* (RA-GHN), a novel framework that systematically unifies graph-based pairwise modeling with biophysical hypergraphs to capture high-order co-operative effects. Specifically, we construct a heterogeneous hypergraph representation featuring three complementary hyperedge types: epitope–paratope interaction patches, spatial structural clusters, and high-degree hub structures, enabling a comprehensive encoding of interface physics. Architecturally, RA-GHN adopts a dual-pathway design that synergistically integrates a *Relation-Aware Equivariant Graph Network* (RA-EGN) with an *Equivariant Hypergraph Encoder*, strictly preserving *SE*(3)-equivariance throughout the sequence–structure co-design process. Furthermore, to mitigate *folding hallucination* and sequence overfitting commonly observed in generative models, we introduce the *Structure–Sequence Decoupling Index* (SSDI), which employs a harmonic mean to non-linearly balance structural physical reliability with sequence novelty. Extensive experiments demonstrate that RA-GHN significantly outperforms state-of-the-art baselines across standard benchmarks and affinity optimization tasks, validating that the fusion of high-order geometric modeling with rigorous evaluation criteria enables the generation of physically feasible and functionally innovative antibodies.

**Highlights:**

- RA-GHN co-designs antibody sequence and structure via hypergraphs.
- Biophysical hyperedges capture high-order effects at binding interfaces.
- Dual equivariant pathways synergistically guarantee local geometry and global shape.
- SSDI metric addresses folding hallucination in antibody evaluation.

## 1. Introduction

Antibodies, as proteins of the immune system, safeguard the host by selectively attaching to antigens, such as viruses and bacteria, providing protection [1]. The interaction between antibodies and antigens is primarily governed by specific regions within the antibodies called complementarity-determining regions (CDRs) [2]. As shown in Fig. 1, it has six hypervariable-loop regions (H1, H2, H3, L1, L2, L3) that are used to recognize and bind an extremely diverse repertoire of antigens. Therefore, in order to create medicinal antibodies capable of effectively targeting specific antigens, designing CDRs is the most crucial step [3, 4].

**Figure 1.**
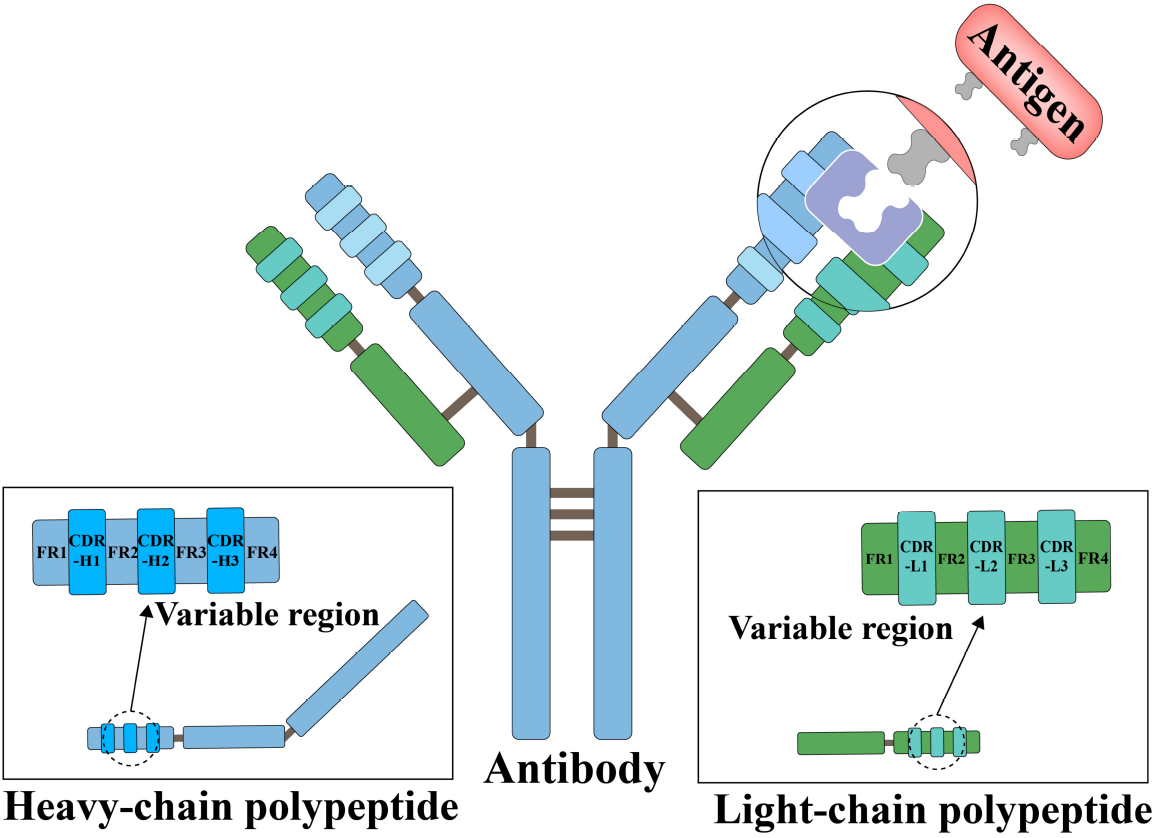
Antibody-antigen complex structure and CDR structure. Antibodies consist of heavy chains and light chains, with each light chain comprising a variable domain (VH/VL) and multiple constant domains. The variable domain is further subdivided into four framework regions (FRs) and three complementarity-determining regions (CDRs).x

CDR diversity is achieved through two key processes: V(D)J gene recombination prior to antigen exposure and somatic hypermutation occurring in B cells after antigen exposure [5]. In biological experiments, the diversification of CDR regions is primarily achieved through techniques such as random mutagenesis, targeted mutagenesis, and chain shuffling. Subsequently, libraries obtained from these diversification methods are screened using display technologies such as yeast or phage display to evaluate their expression and binding characteristics [6]. However, a CDR sequence containing *L* amino acids has up to 20^*L*^ possible protein sequences, making it impractical to test all potential sequences experimentally, and the small solution space that satisfies the desired constraints of binding affinity, stability, and synthesizability [2, 7]. Therefore, computer-aided antibody design can effectively address this challenge.

Traditional computational approaches rely on sampling protein sequences and structures from complex theoretical energy functions (physics-based) or statistical (knowledge-based) energies [8, 9, 10, 11, 12]. However, these methods typically suffer from long computational times and are susceptible to being trapped in local optima. Furthermore, no force field comprehensively captures the intricate interactions inherent in biological molecules. In recent years, with the rapid advancement of deep generative models, several approaches have emerged for antibody design and protein design [2, 6, 13, 14, 15, 16]. Compared to traditional algorithms, deep generative models can directly capture the higher-order interactions between amino acids in antibodies and antigens, resulting in more effective antibody generation [17]. However, there is still a significant gap that needs to be addressed in order for generative models to become practical for antibody design.

Early studies of antibody design focused mainly on the generation of CDR sequences without simultaneously considering their corresponding structural information[13, 14, 18]. Subsequently, Jin et al.[19] proposed the RefineGNN method, which achieves joint design (co-design) of CDR sequences and structures, providing a new perspective for the coordinated optimization of antibody structure and function. The subsequent work HSRN[20] integrates the design and docking of antigen-binding antibodies within a hierarchical refinement framework. Building upon this, DiffAb[2] utilizes diffusion models [21] to iteratively generate CDR sequences, C*α* coordinates, and orientation information. Similarly, AbDiffuser[22] incorporates family-specific priors into the diffusion process, significantly enhancing both generation efficiency and quality compared to DiffAb. To improve computational efficiency, MEAN[23] replaces autoregressive decoding with progressive full-shot decoding and incorporates light chain information as a conditional input for CDR generation. This approach was later extended to dyMEAN[24], enabling end-to-end full-atom antibody design. A recent study RAAD[25] models each antigen–antibody complex as an Attributed Heterogeneous Graph (AHG), defining multiple edge types to enrich residue sequence and structural context. It performs message passing for each relation separately and integrates the results into a final representation. By taking the entire antigen as input and modeling the dynamics of the antigen-antibody interaction as an optimization problem of the edge relationship, RAAD significantly improves the sequence–structure co-design of antibodies.

However, as the conditional contexts become increasingly complex, such models still face significant challenges in effectively capturing the intricate structural and sequential interactions between CDRs and their contexts. The fundamental limitation lies in their insufficient geometric modeling capability—existing methods rely solely on local residues or pairwise relationships, making it difficult to fully capture multi-body interactions and global structural dependencies in antibody molecules, thereby limiting the representation of complex geometric constraints and sequence–structure cooperative effects.

To address the limitations of existing methods in antibody sequence–structure co-design, we propose a novel model—Relation-Aware Graph–Hypergraph Networks (RA-GHN). Unlike traditional approaches that rely solely on local residues or pairwise relationships, RA-GHN leverages the strengths of both graphs and hypergraphs to simultaneously capture second-order and higher-order multi-residue interactions within antibody molecules. Through this joint modeling strategy, the model can not only capture local dependencies between sequence and structure but also explicitly represent global geometric constraints and complex spatial conformations, providing a more comprehensive reflection of the functional and structural cooperativity of antibodies.

During message passing, RA-GHN performs relation-aware message passing for different types of edge relations and integrates contextual information from multiple relations into a final representation enriched with higher-order structural dependencies. This approach effectively overcomes the shortcomings of traditional methods in modeling long-sequence CDRs, complex geometric constraints, and multi-residue cooperative effects, significantly enhancing the representation of antibody sequence–structure interactions and providing a solid foundation for high-precision antibody design and optimization.

Specifically, we decompose antibody CDR design into two core stages: (i) Graph–Hypergraph Geometric Modeling and (ii) Sequence–Structure Co-Learning. In the geometric modeling stage, RA-GHN leverages graph structures to capture second-order residue interactions and uses hypergraphs to explicitly represent higher-order multi-residue interactions, thereby comprehensively modeling the complex spatial geometry and global constraints within antibody molecules. It is important to note that here, “complex spatial geometry” does not refer to protein conformational dynamics, but rather to the static geometric relationships and interaction patterns among residues within the antibody and the antigen–antibody complex. This approach simultaneously captures local dependencies within the residues of the antigen–antibody complex as well as global interactions between different components, providing a solid geometric foundation for sequence–structure co-design. In the sequence–structure co-learning stage, we introduce a relation-aware message passing mechanism that propagates information for different types of edge relations separately and integrates the contextual information into the final representation, enabling complementary optimization of sequence and structure features. This strategy avoids the need for computationally expensive explicit iterative refinement while effectively enhancing the model’s ability to represent complex multi-residue interactions. We evaluated RA-GHN on three challenging tasks: sequence and structure modeling, antigen-binding CDR design, and binding affinity optimization. Experimental results demonstrate that our model outperforms existing state-of-the-art methods across all tasks, validating its effectiveness and utility in antibody sequence–structure co-design.

The main contributions of this work are summarized as follows:

- We propose the *Relation-Aware Graph-Hypergraph Network* (RA-GHN), a novel framework that systematically unifies graph-based pairwise residue interactions with hypergraph-based high-order cooperative effects. To the best of our knowledge, this is the first *SE*(3)-equivariant architecture to seamlessly integrate these dual geometric modalities for the simultaneous optimization of antibody CDR sequences and structures.
- We construct a biophysically grounded hypergraph representation comprising three complementary classes of hyperedges: epitope–paratope interaction patches, spatial structural clusters, and high-degree interaction hubs. This multi-view formulation enables the explicit modeling of complex multi-body dependencies and global geometric constraints that are critical for shaping the protein–protein interaction interface.
- We engineer a dual-pathway fusion mechanism that processes structural information in parallel via a *Relation-Aware Equivariant Graph Network* (RA-EGN) and an *Equivariant Hypergraph Encoder*. This design synergistically balances local bond geometry with global shape consistency while strictly guaranteeing rotational and translational equivariance throughout the generation process.
- We introduce the *Structure–Sequence Decoupling Index* (SSDI) to address the limitations of existing evaluation metrics, specifically the risks of “folding hallucination” and sequence overfitting. By employing a harmonic mean to nonlinearly couple structural reliability with sequence novelty, SSDI provides a robust and discriminative criterion for identifying antibody designs that are both physically feasible and functionally innovative.

## 2. Preliminary

As illustrated in Fig. 1, antibodies are Y-shaped proteins composed of heavy and light chains, each containing a variable domain (*V*_*H*_ /*V*_*L*_). These domains feature hypervariable Complementarity-Determining Regions (CDRs) flanked by framework regions. Since the heavy-chain CDRs (H1–H3) are the primary determinants of antigen-binding specificity and affinity, this study focuses specifically on their computational design and optimization.

To capture the multi-scale interactions inherent in antigen–antibody binding, we propose a *Hybrid Topology* comprising two complementary components: 1) a *Pairwise Relational Graph* (*G*) for modeling pairwise relations; 2) a *Biophysical Hypergraph* (*H*)for capturing high-order cooperative effects.

### 2.1. Pairwise Relational Graph

We adopt the heterogeneous graph formulation from RAAD [25] to represent the antigen–antibody complex **as a collection of pairwise residue interactions**. The complex is modeled as *G* = (*V*, ℰ), where nodes *V* represent residues endowed with geometric and semantic features, and edges ℰ capture multi-scale interactions between **node pairs**.

#### Featurization and Relations

Input features are designed to be invariant to rigid body transformations. Node features aggregate residue identity, sinusoidal positional encodings, and local backbone geometry (including angles and coordinates). Edge features capture relative geometric information, specifically RBF-encoded distances and quaternion-based relative orientations. To systematically model the hierarchy of interactions, edges are categorized into eight relation types covering sequential connectivity, intra/inter-chain spatial proximity, and global context connections [25]. This rigorous graph construction serves as the foundation for our subsequent hypergraph evolution.

### 2.2. Biophysical Hypergraph

To model the intricate, high-order cooperative effects inherent in antigen–antibody complexes—which fundamentally extend beyond simple pairwise interactions—we introduce a biophysically grounded hypergraph representation *H* = (*V*, ℰ_hyper_). In traditional graph-based molecular representations, edges are restricted to connecting exactly two nodes, implicitly assuming that thermodynamic and geometric contributions are purely additive. However, at protein–protein interfaces, binding affinity and structural specificity are frequently dictated by synergistic many-body interactions, such as cooperative hydrogen bond networks, dense multi-residue hydrophobic packing, and coordinated electrostatic environments. Unlike standard graphs, a hyperedge *e* ∈ ℰ_hyper_ can concurrently encompass an arbitrary number of residue nodes *v* ∈ ℰ. This mathematical property provides a natural and highly expressive formulation for the many-body cooperativity prevalent at these complex binding interfaces.

To systematically construct this high-order topology, we design a biophysics-aware hyperedge constructor, denoted as Φ_hyper_. Rather than relying solely on naive global distance cutoffs, Φ_hyper_ generates semantically and physically meaningful hyperedges tailored to the fundamental biophysical principles of antigen–antibody recognition. Specifically, we constructed three types of complementary hyperedges, each targeting a different aspect of the structure-energy interaction:we formulate three complementary classes of hyperedges, each addressing a distinct aspect of the structural and energetic interplay:

- **Epitope–Paratope Interaction Patches:** Captures cross-chain spatial packing and shape complementarity directly across the binding boundary, modeling how local groups of residues function as integrated units to cooperatively accommodate specific antigen features.
- **Spatial Structural Clusters:** Encapsulates dense, intra-chain topological packing. This captures the spatial aggregation necessary to stabilize local folding units and preserve the anchoring architecture of highly flexible Complementarity-Determining Regions (CDRs).
- **High-Degree Interaction Hubs:** Identifies topologically central energetic hotspots using a graph-theoretic approach, establishing star-topology structures that efficiently route geometric and feature signals from critical binding residues to their localized neighborhoods.

By seamlessly integrating these three complementary hyperedge types, our representation mathematically guarantees that the model perceives the antibody–antigen complex not merely as a collection of isolated atomic contacts, but as a tightly coupled, heterogeneous biochemical system. The detailed construction algorithms and formulations for each hyperedge class are elaborated below.

#### Epitope–Paratope Interaction Patches

The high-affinity and highly specific recognition between antibodies and antigens is rarely governed by isolated, independent residue–residue contacts. Instead, it relies heavily on the synergistic formation of local surface patches composed of multiple spatially proximal residues acting in concert. These interaction patches exhibit a profound degree of multi-dimensional complementarity—encompassing topological geometric shape (e.g., knob-into-hole packing), electrostatic potential pairing, and optimal hydrophobic/hydrophilic matching. Consequently, they constitute the fundamental functional and energetic units of the antigen–antibody binding interface. Modeling these molecular interactions solely at the level of pairwise residue contacts fundamentally limits the representational capacity of the network, as it is often insufficient to capture the highly cooperative nature of interface complementarity, such as multi-centered hydrogen bond networks or distributed hydrophobic interfaces.

To explicitly model this patch-level cooperativity at the antigen–antibody interface, we introduce cross-chain interaction patches (epitope–paratope patches) and formally represent them as hyperedges. Let *V*_*Ab*_ and *V*_*Ag*_ denote the distinct sets of interface residues on the antibody (comprising both the heavy and light chain variable domains) and the target antigen, respectively. For each core antibody interface residue *u* ∈ *V*_*Ab*_, we construct a bi-partite, patch-based hyperedge 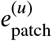 structurally centered at *u*. This hyperedge simultaneously aggregates the target antigen residues that are spatially proximal to *u* across the binding interface, along with its structurally adjacent antibody residues. This dual-aggregation strategy successfully bridges the intra-chain structural scaffold with the inter-chain binding targets, thereby forming a cohesive local cross-chain interaction unit:

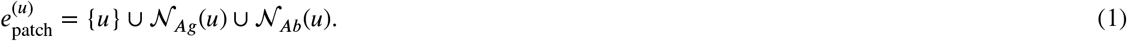

The cross-chain antigen neighborhood is mathematically defined as:

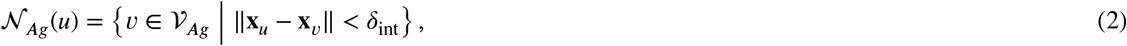

where **x**_*u*_ and **x**_*v*_ denote the three-dimensional atomic coordinates (e.g., *C*_*α*_ or side-chain centroids) of residues *u* and respectively. The spatial distance threshold is meticulously set to δ_int_ = 6.0 Å. This specific cutoff corresponds to the typical biophysical range necessary to capture diverse non-covalent antigen–antibody interfacial contacts, effectively encompassing direct van der Waals interactions, strong electrostatic salt bridges, and potentially water-mediated hydrogen bonds, while naturally filtering out distant, non-interacting residues.

In dense interfacial regions, an unrestricted neighborhood search could lead to excessive patch sizes, which implicitly introduces redundant structural noise and exacerbates the risk of over-smoothing during message passing. To prevent this while strictly preserving local geometric coherence and computational tractability, we impose a hard upper bound on the number of participating nodes within each patch-based hyperedge. The maximum capacity is constrained to *K*_patch_ (empirically set to *K*_patch_ = 5 in this study). When the number of candidate neighboring residues exceeds this predefined threshold, the nodes are ranked by their Euclidean distance to the center *u*, and only the spatially closest residues are retained. This truncation ensures that the hyperedge precisely captures the most energetically dominant and sterically relevant contacts.

Through this biophysically motivated construction, the model intrinsically treats the local geometric configuration, steric constraints, and chemical microenvironment of the binding interface as a single integrated functional unit, fundamentally departing from the limitations of independent pairwise representations. This patch-level modeling strategy substantially enhances the network’s capacity to explicitly capture holistic interface complementarity, synergistic energetic effects, and unique local geometric fingerprints (e.g., structural motifs distinct to specific paratope– epitope pairings). Consequently, it provides a significantly richer and more biophysically meaningful structural basis for downstream high-order information propagation, ultimately improving the discrimination of high-affinity binders from structural decoys.

#### Spatial Structural Clusters

To rigorously maintain the stability of a protein’s tertiary structure, spatially proximal residues are driven by thermodynamic imperatives—such as the burial of hydrophobic side chains—to cooperatively pack into compact structural clusters. Such spatial structural clusters play a paramount role in protein folding: they severely restrict the conformational entropy of both the backbone and side chains, stabilize local folding units (e.g., *α*- helices or dense β-sheet cores), and preserve the overall topological organization of the macromolecule. In the context of antibodies, although the complementarity-determining regions (CDRs) inherently exhibit pronounced structural flexibility to accommodate diverse antigens, their stable and functional conformations (i.e., canonical structures) remain critically dependent on the ordered aggregation and rigid structural scaffolding provided by surrounding framework residues in three-dimensional space.

To explicitly characterize these intra-chain spatial aggregation effects, we employ a geometry-based greedy clustering algorithm to identify dense spatial structural clusters within the proteins and formally model them as hyperedges. To rigorously circumvent cross-chain topological interference and preserve the distinct conformational identity of each molecule prior to complex formation, the algorithm is executed independently on each distinct protein segment (i.e., the antibody heavy chain, light chain, and the target antigen). Let *U* denote the dynamic pool of residue nodes that have not yet been assigned to any structural cluster. The algorithm incrementally constructs these spatial clusters through an iterative procedure, as illustrated in Algorithm 1.

##### Algorithm 1

Construction of Spatial Structural Cluster Hyperedges

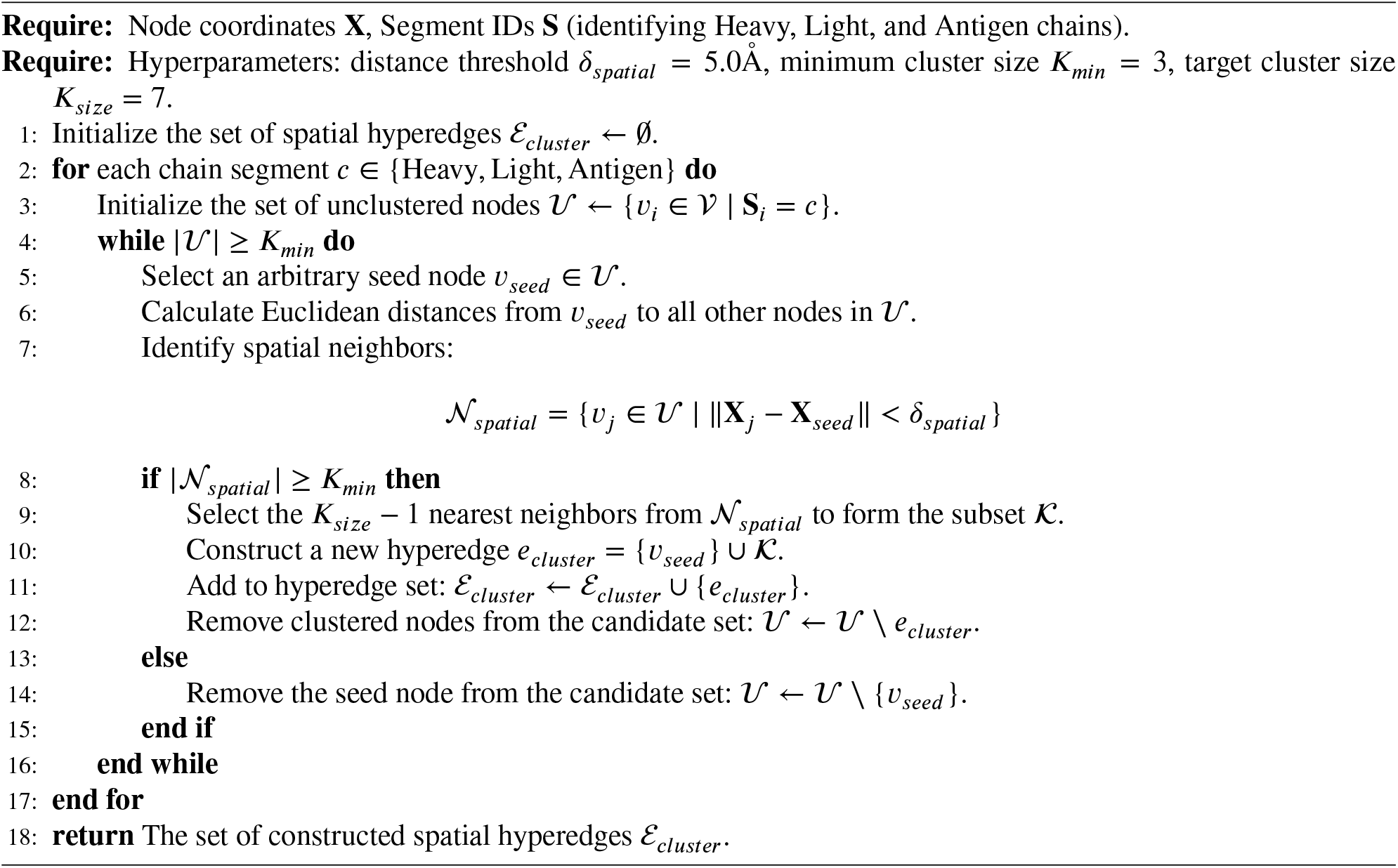

The algorithm requires the 3D residue coordinates **X** and chain segment identifiers **S** as inputs. As outlined in the pseudocode, the procedure iteratively processes the set of unclustered residues *U* utilizing a greedy selection strategy (Lines 5–16) to discover candidate spatial motifs. In each iteration, a seed residue is randomly sampled from *U*, and its spatial neighborhood is delineated by a Euclidean distance threshold δ_spatial_. This critical threshold is biophysically calibrated to capture the typical scale of tight residue packing, effectively defining the first coordination shell of intra-protein contacts. It is particularly adept at detecting highly cooperative interactions, such as deep hydrophobic cores or internal stabilizing hydrogen-bond networks anchoring the CDR loops.

The biophysical validity of each candidate cluster is then rigorously assessed against a minimum density requirement, *K*_min_ (Line 9). This density filter ensures that only sufficiently populated, energetically significant clusters are retained, effectively discarding sparse, highly solvent-exposed, or transiently interacting residues that would otherwise introduce structural noise into the graph. Once a candidate cluster satisfies this density criterion, a structural hyperedge *e*_cluster_ is instantiated. To prevent the formation of excessively massive hyperedges—which could obscure local geometric details and lead to signal over-smoothing across the deep neural network—the cluster size is strictly bounded. Specifically, the hyperedge incorporates the seed residue along with its spatially closest unassigned neighbors, up to a maximum capacity of *K*_size_. All residues assimilated into this newly formed hyperedge are subsequently removed from the unassigned pool *U* to guarantee mutually exclusive cluster assignments.

This iterative extraction process continues exhaustively until no further valid clusters can be formed from the remaining nodes in *U*. The final output of the algorithm is the set of spatial cluster hyperedges ℰ _cluster_, which encodes a robust topological map of compact structural motifs formed by spatially adjacent residues. These intra-chain hyperedges are subsequently merged with other hyperedge modalities (e.g., cross-chain interface patches and interaction hubs) to construct a unified, heterogeneous structural representation. By capturing both strict local geometric constraints and cooperative residue packing, these clusters serve as a physically meaningful structural prior, strongly anchoring the downstream high-order information propagation and geometric representation learning.

#### High-Degree Interaction Hubs

In the thermodynamic landscape of antigen–antibody binding interfaces, the distribution of binding free energy is notoriously highly localized. A small subset of critical residues—often referred to as binding hotspots—functions as the primary energetic anchors, contributing disproportionately to the overall binding affinity while surrounded by energetically neutral peripheral residues (akin to the biophysical “O-ring” theory). These hotspot residues are not only strategically located at key spatial positions but also exhibit exceptionally high centrality within the network topology; the density of non-covalent interactions among their immediate neighbors significantly exceeds the interface average. To explicitly endow the neural network with the ability to focus its receptive field on these critical regions, we introduce a topology-driven hub hyperedge construction strategy. This approach structurally incorporates high-degree residues and their local microenvironments into unified, attention-demanding hyperedge representations.

Specifically, we first construct an underlying interface interaction graph *G*_int_ = (*V*_int_, ℰ_int_), where the node set *V*_int_ represents all identified interface residues, and the edge set ℰ_int_ encodes basic pairwise residue–residue interactions defined by spatial proximity. For each node *v* ∈ ℰ_int_, we compute its topological degree deg(*v*) to quantitatively assess its network connectivity and centrality within the interfacial environment. Because high topological connectivity strongly correlates with energetic hotspot propensity, we select the top-*K* nodes possessing the highest degrees to populate the hub set *H*_nodes_ (with *K* = 10 in this study, ensuring a consistent computational focus across complexes of varying sizes).

For each identified hub node *h* ∈ *H*_nodes_, we construct a targeted, star-topology hyperedge 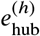. Instead of indiscriminately grouping all neighbors, we rigorously define this hyperedge by connecting the central hub node specifically to its *M* nearest topological neighbors within the interface graph, thereby forming a highly condensed local interaction unit:

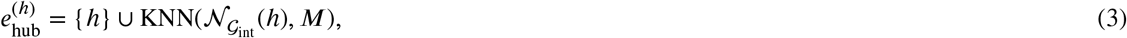

where 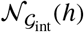 denotes the comprehensive set of neighbors of node *h* in the base interface graph, and *M* specifies the maximum capacity of neighbor nodes (strictly set to *M* = 6 in this study to filter out distant, low-impact contacts). This mathematical construction imposes a strict star-topology structure: the central energetic hub radiates outward to its immediate structural scaffolding. This topology acts as an efficient “information highway,” seamlessly routing geometric and biochemical signals from the critical hub directly to its surrounding microenvironment during message passing.

By formally introducing these hub hyperedges, the model is mathematically compelled to distinguish between the core energetic engines of the interface (hub residues and their densely packed neighborhoods) and the structurally supportive but energetically peripheral regions. Within the paradigm of graph neural network-based information propagation, hub hyperedges effectively prevent critical hotspot signals from being diluted or “drowned out” by the majority of less important residues during global feature aggregation. Instead, they amplify the structural contribution of key residues across multiple deep network layers, significantly enhancing the model’s sensitivity and predictive accuracy for binding free energy estimation and hotspot identification. Furthermore, this topological approach is intrinsically orthogonal and naturally compatible with our geometry-based hyperedges (i.e., patch-based units and spatial structural clusters). Together, they coalesce into a unified, heterogeneous hypergraph architecture that simultaneously captures localized geometric constraints, chemical complementarity, and global topological centrality, thereby establishing an exceptionally rich and biophysically coherent representation for the high-order structural modeling of antigen–antibody interactions.

#### Hyperedge Merging and Optimization

In the modeling of antibody–antigen interfaces, three types of hyperedges are constructed: cross-chain interaction patches (ℰ_*patch*_), spatial structural clusters (ℰ_*cluster*_), and high-degree interaction hubs (ℰ). While each hyperedge type captures different levels of structural or functional information, processing them separately in the model can lead to computational inefficiency and inconsistent feature spaces. To improve computational efficiency and unify feature representation, we introduce a hyperedge merging strategy that consolidates all hyperedges into a single sparse incidence matrix.

It is worth emphasizing that although all hyperedges are concatenated into a single sparse incidence matrix, this operation is merely a computational reorganization and does not imply any semantic conflation among different hyperedge types. The distinction between patch-based interfaces, spatial clusters, and hub residues is implicitly preserved by their unique node composition patterns and the distinct physicochemical features of the constituent residues. During the subsequent message passing process (Eq. (9)), the model learns to assign different importance and update patterns to different hyperedges based on their structural contexts, rather than treating them as identical objects. In other words, the unified matrix provides a convenient and efficient computational scaffold, while the semantic heterogeneity is automatically captured by the learning process.

Formally, let the complete set of constructed hyperedges be:

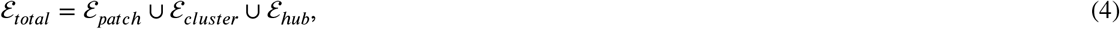

where ℰ_*patch*_, ℰ_*cluster*_, and ℰ_*hub*_ represent patch-based hyperedges, spatial cluster hyperedges, and hub hyperedges, respectively. During the merging process, hyperedge indices are re-ordered to form a contiguous numbering, resulting in a sparse incidence matrix:

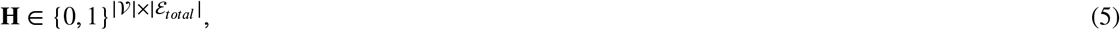

where each column corresponds to a unique hyperedge and the matrix entries are defined as:

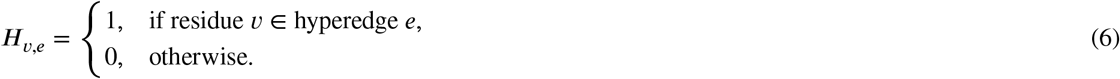

The unified incidence matrix standardizes the input format for parallel computation; however, the semantic diversity of different hyperedge categories is preserved through the distinct connectivity patterns and node feature distributions associated with each type, ensuring that the model can differentiate them during parameter learning.

This unified sparse matrix simultaneously encodes the relationships between residues and high-order interaction units. The sparsity of **H** reduces both memory and computational costs, which is especially important for large-scale interface modeling.

Moreover, the unified incidence matrix provides a convenient input structure for Equivariant Hypergraph GNNs. During training or inference, the network can process all hyperedge types in parallel, without separate loops for patches, spatial clusters, or hub hyperedges. This not only simplifies the data pipeline but also ensures that different types of high-order information are represented in a shared feature space. Consequently, the model can integrate local geometric features, residue packing patterns, and topological information of key binding sites within a single high-order graph representation, providing a unified and efficient computational framework for interface characterization, binding affinity prediction, and structure-sensitive information propagation.

## 3. Model Architecture

### 3.1. Overview

As illustrated in Fig. 2, the proposed novel model, Relation-Aware Graph-Hypergraph Network for Antibody Sequence–Structure Co-Design (RA-GHN), consists of multiple stacked relation-aware graph–hypergraph layers, which integrate equivariant pairwise geometric message passing with hyperedge-based higher-order interaction modeling. By jointly reasoning over residue-level graphs and biologically motivated hypergraphs, RA-GHN enables coherent generation of sequences and structures for antibody CDRs.

**Figure 2.**
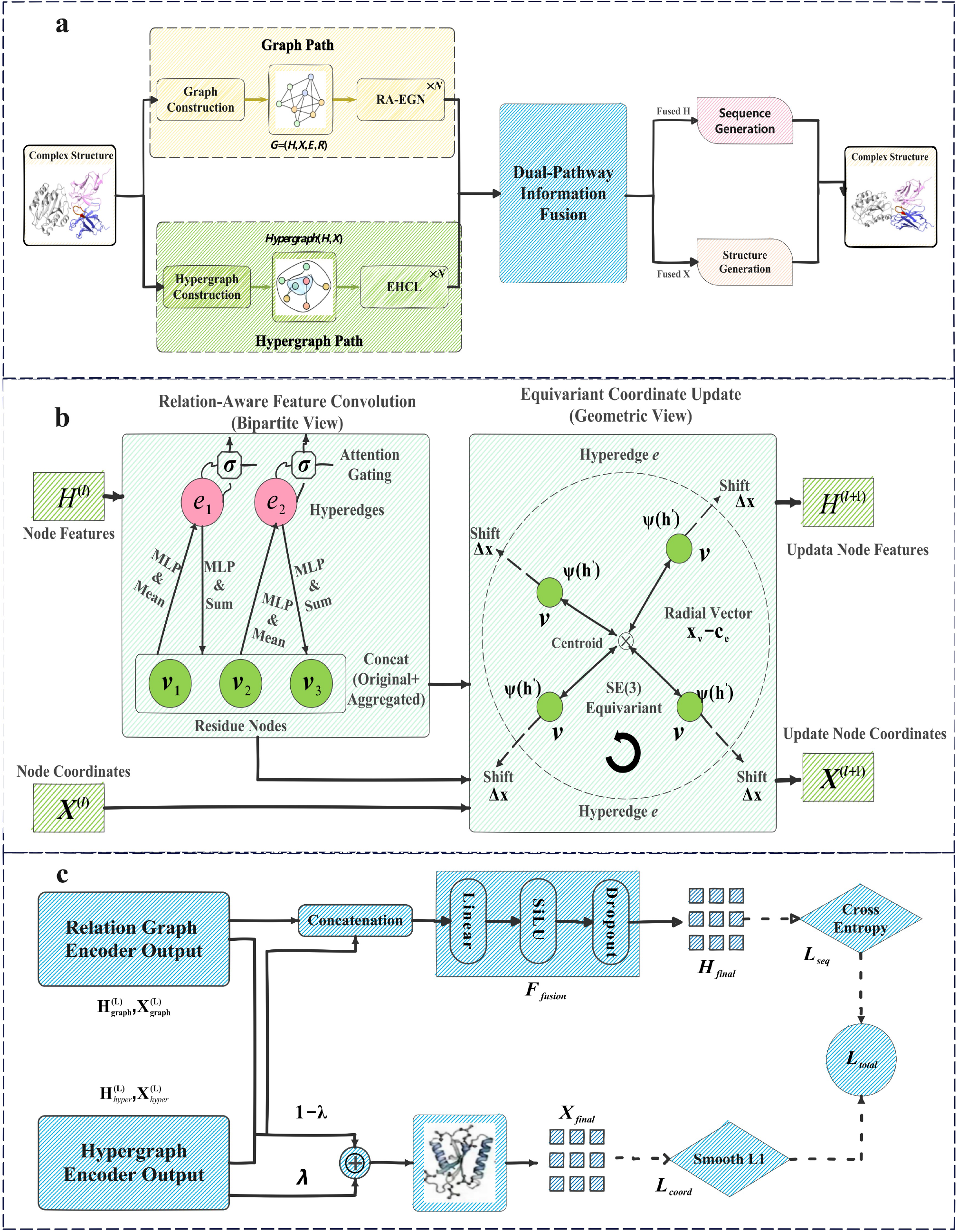
The overview of Relation-Aware Graph-Hypergraph Network(RA-GHN). (a) The overall framework of RA-GHN. (b) The schematic of Equivariant Hypergraph Convolutional Layers (EHCL). (c) The schematic of the Dual-Pathway Fusion and Prediction Module.

### 3.2. Pairwise Interaction Modeling via Relation-Aware EGNN

To capture the intricate local geometric constraints and fine-grained pairwise interactions—such as hydrogen bonds, Van der Waals forces, and electrostatic contacts—within the complex, we employ the *Relation-Aware Equivariant Graph Network* (RA-EGN) [25] as the backbone of our graph encoder. Unlike standard graph neural networks that operate solely on topological connectivity, RA-EGN iteratively refines node features, edge attributes, and atomic coordinates within a unified spatial framework. Crucially, it achieves this while strictly preserving *SE*(3)-equivariance, ensuring that the modeled spatial dynamics and energy landscapes remain completely invariant to the arbitrary choice of the global coordinate reference frame.

Formally, given a relation graph *G*, the encoder updates representations at layer *l* through a relation-sensitive spatial message passing mechanism. For a target residue *i* and its neighbor *j* connected by a specific relation type *r* (which inherently encodes sequence separation or specific chemical bond properties), the geometric message 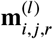 is computed as:

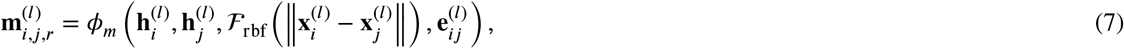

where ϕ_*m*_ denotes a shared multilayer perceptron (MLP) acting as the message function. The term *F*_rbf_ (⋅) smoothly projects the continuous pairwise Euclidean distances into a high-dimensional continuous feature space using radial basis functions, enabling the network to sensitively capture distance-dependent physical interactions. Additionally, 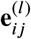 incorporates intermediate edge representations that continuously evolve to capture dynamic inter-residue relationships.

The aggregated messages are subsequently utilized to jointly update both the node features **h**_*i*_ and the edge features **e**_*ij*_, effectively propagating local structural context throughout the graph. More importantly, to physically adapt the 3D conformation, the atomic coordinates are updated in a vector-valued manner:

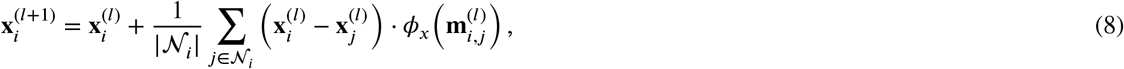

where *N*_*i*_ denotes the set of spatial neighbors for node *i*, and ϕ (⋅) is a learnable scalar mapping derived from the aggregate^*i*^d messages. Intuitively, this coordinate update acts as a neural simulated force field: ϕ_x_ predicts the magnitude and direction (attractive or repulsive) of the interaction, and the spatial displacement is strictly constrained along the relative radial vector 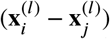. This vector-based formulation rigorously guarantees *SE*(3)-equivariance for spatial dynamics while prevent^*i*^ing un^*j*^physical coordinate shifts.

Overall, this relation-aware pairwise module effectively models the immediate, micro-level physical dependencies between adjacent residue pairs. By establishing a robust and physically grounded local structural embedding, it seamlessly complements the high-order, macroscopic topological representations extracted by the hypergraph encoder, resulting in a comprehensive hierarchical modeling of the multi-chain complex.

### 3.3. Equivariant Hypergraph Encoder

To efficiently capture high-order interactions while strictly preserving *SE*(3)-equivariance, we propose an *Equivariant Hypergraph Encoder* composed of *L* stacked convolutional layers. Unlike clique-expansion methods, we adopt a *bipartite message passing* mechanism that directly exchanges information between nodes and hyperedges. The proposed encoder consists of *L* stacked *Equivariant Hypergraph Convolutional Layers*(EHCL). A schematic of EHCL is shown in Fig. 2(b).

#### Relation-Aware Feature Convolution

To effectively capture and fuse the diverse biophysical contexts inherent in complex molecular environments, we introduce an attention-gated hypergraph convolution mechanism. Standard message-passing algorithms often struggle to differentiate between critical structural motifs and coincidental spatial proximities. To overcome this, our feature update at layer *l* proceeds through three rigorously defined logical steps:

i. *Node-to-Hyperedge Aggregation:* Each hyperedge *e* functions as a high-order receptive field, aggregating localized biochemical and geometric information from its constituent nodes *V*_*e*_. We define the high-order hyperedge representation as:

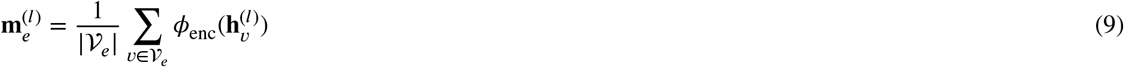

where ϕ_enc_ is a neural network module (e.g., a multilayer perceptron) that projects individual residue embeddings into a shared latent semantic space, effectively capturing the collective biochemical signature of the multi-body interaction unit.
ii. *Attention Gating:* In distance-based geometric graphs, sparse or transient neighbors can introduce significant structural noise. To dynamically filter this noise and isolate functionally relevant interactions, we introduce a learnable scalar attention mechanism:

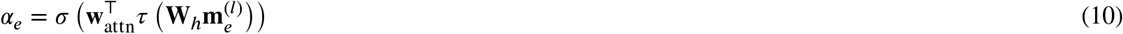 This gating function evaluates the structural and energetic relevance of the hyperedge, yielding a continuous weight *α*_e_. The reweighted, noise-filtered hyperedge feature is then computed as 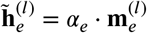
iii. *Propagation & Update:* Finally, these refined, high-order contextual messages must be propagated back to the individual constituent nodes. A fundamental challenge in deep geometric networks is the “over-smoothing” phenomenon, where repeated message passing causes node features to lose their distinct topological identities. To explicitly mitigate this, we concatenate the newly aggregated macroscopic message with the node’s original feature representation prior to the final update:

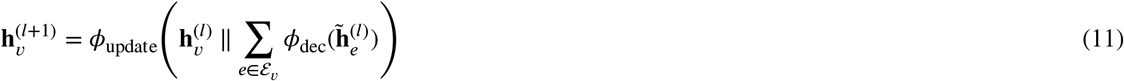

This residual-like concatenation ensures that each residue strictly preserves its unique biochemical identity while seamlessly integrating broader neighborhood awareness.

#### Centroid-Based Equivariant Coordinate Update

To translate abstract learned feature representations into physically meaningful 3D spatial conformations, we propose a geometric coordinate update strategy governed by *hyperedge centroids*. The core intuition is to conceptualize the coordinate update as a dynamic, neural-simulated radial force field exerted by the local macro-structural context.

First, we determine the spatial focal point of the multi-body interaction by computing the geometric centroid of each hyperedge:

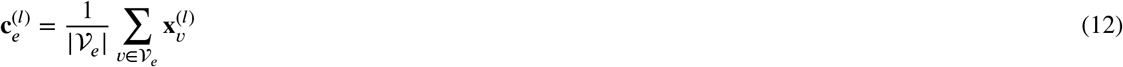

The spatial displacement for a given node *v* is then derived as the weighted sum of relative position vectors, which point from the respective hyperedge centroids directly to the target node:

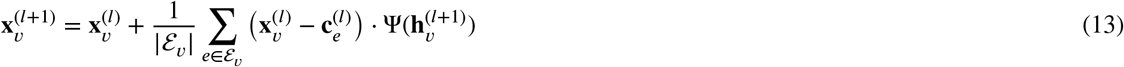

where Ψ(⋅) represents a neural mapping function that predicts the magnitude and directional sign (attractive or repulsive) of the spatial displacement based on invariant node features. Because the relative directional vector (**x**_*v*_ −**c**_*e*_) rotates and translates perfectly equivariantly with any transformation of the global reference frame, and the scaling weight Ψ is strictly derived from invariant scalars, this vector-based aggregation mathematically guarantees exact *SE*(3)-equivariance throughout the network’s spatial dynamics.

Crucially, the biophysical validity of this radial formulation is deeply grounded in our model’s exclusive focus on backbone-only generation. By temporarily stripping away highly flexible, bulky, and structurally diverse amino acid side chains, the remaining backbone nodes function as coarse-grained topological control points. In this specific context, pulling nodes toward a local geometric centroid does not constitute a forceful, microscopic atomic compression that would typically induce severe steric clashes or bond-angle distortions in an all-atom model. Rather, it geometrically emulates the macroscopic thermodynamic phenomenon of “hydrophobic collapse” observed during the early stages of protein folding—efficiently guiding dispersed secondary structural elements to aggregate toward a local center to form a compact, globally consistent core. While this macroscopic neural force field prioritizes the rapid convergence of the global topology over strict, localized microscopic rigidity, the resulting backbone scaffolds provide highly tolerant, shape-accurate 3D templates. Any minor deviations from ideal local stereochemistry can be seamlessly restored during downstream sequence design (inverse folding) and physical force-field relaxation.

### 3.4. Dual-Pathway Information Fusion

To seamlessly integrate the fine-grained, pairwise structural constraints derived from the relation graph with the high-order, multi-body cooperative effects captured by the hypergraph, we introduce a robust adaptive fusion module. In complex biomolecular systems, local stereochemistry and global topological packing are deeply interdependent; thus, this module is specifically designed to project both heterogeneous representations into a unified, physically coherent prediction space. A detailed schematic of the Dual-Pathway Fusion and Prediction Module is illustrated in Fig. 2(c).

#### Adaptive Feature Fusion

To capture complementary local and global semantics without premature information bottlenecking, we first concatenate the final layer output features from both pathways, denoted as 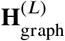 and 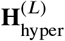. This concatenation explicitly preserves the orthogonal geometric and biochemical features extracted by each distinct branch. A nonlinear fusion network, *F*_fusion_, then maps this combined high-dimensional representation directly to the amino acid logits:

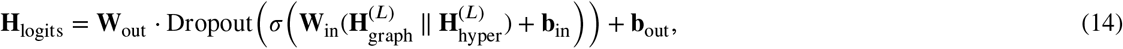

where σ(⋅) denotes the SiLU (Swish) activation function, systematically chosen for its smooth, non-monotonic properties that facilitate stable gradient flow during deep network optimization. The inclusion of the Dropout layer serves as a critical structural regularizer, effectively preventing the model from over-relying on a single informational pathway (e.g., overfitting strictly to pairwise distances while neglecting hypergraph topology). Ultimately, this network adaptively balances multi-scale structural information to accurately predict residue-type probabilities for downstream sequence design.

#### Weighted Coordinate Aggregation

Beyond abstract feature representations, predicting physically viable 3D coordinates requires a delicate equilibrium between microscopic local bond fidelity and macroscopic shape consistency. To achieve this, we reconcile the spatial predictions from both pathways via a weighted ensemble strategy. The final refined atomic coordinates are computed as a linear interpolation:

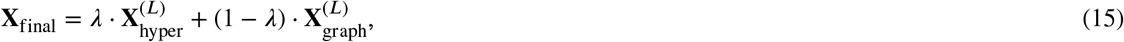

where the balancing coefficient λ ∈ [0, 1] dynamically modulates the structural trade-off. Specifically, it scales the balance between global structural coordination (driven by the hypergraph’s macro-scale topological pull) and local geometric precision (maintained by the graph’s micro-scale distance constraints). By organically blending these spatial predictions, this ensemble mechanism effectively mitigates coordinate noise and spatial jittering. It mathematically guarantees that the generated conformations simultaneously satisfy rigorous local chemical validity (e.g., ideal bond lengths and absent steric clashes) and holistic interface compatibility.

### 3.5. Optimization Objectives

The end-to-end training of the proposed model is jointly guided by the sequence generation task and the structure prediction task. Because the sequence identity and the 3D conformation of highly flexible Complementarity-Determining Regions (CDRs) are intrinsically coupled biophysical properties, co-optimizing them simultaneously ensures that the generated sequences are structurally viable and the predicted backbones are biochemically sound. The overall optimization objective is defined as a weighted sum of the two task-specific losses:

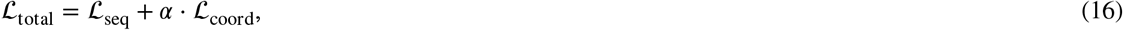

where α is a tunable hyperparameter that dynamically balances the relative contributions of sequence-level and structure-level supervision, ensuring that neither modality dominates the gradient updates during the learning process.

#### Sequence Reconstruction Loss (*L*_seq_)

To stringently supervise the amino acid type prediction strictly within the highly variable masked CDR region *M*, we employ the standard cross-entropy loss. This objective functions as a constrained inverse-folding mechanism, penalizing biochemical deviations from the native sequence based on the extracted spatial geometry. Formally:

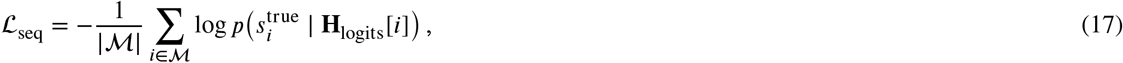

where 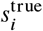 denotes the one-hot encoded ground-truth amino acid identity at sequence position *i*, and **H**_logits_[*i*] represen^*i*^ts the predicted raw logit distribution over the standard amino acid vocabulary. By optimizing this loss, the network learns to interpret the fused dual-pathway structural representations to recover the optimal sequence corresponding to the geometric template.

#### Structure Coordinate Loss (*L*_coord_)

To explicitly supervise the predicted 3D backbone coordinates of the generated CDR residues, we adopt the Smooth L1 (Huber) loss. Unlike the standard Mean Squared Error (L2 loss), which is highly sensitive to significant spatial deviations and can trigger exploding gradients, Smooth L1 provides a critical optimization balance. It behaves identically to an L1 loss for large structural outliers—which are prevalent during the initial modeling phases of highly flexible loops—and seamlessly transitions to an L2 loss for minor local deviations to encourage precise spatial convergence. The loss is computed as:

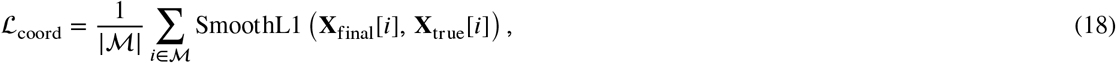

where **X**_final_[*i*] and **X**_true_[*i*] denote the predicted and ground-truth 3D backbone coordinates (e.g., *C*_α_ anchor atoms) of residue *i*, respectively. This robust and continuous constraint mathematically guarantees that the model accurately captures the physical anchoring and macroscopic geometric shape of the CDR loop without overly penalizing the inherent natural flexibility of the backbone.

## 4. Experiment setup

### 4.1. Experimental Design

Extensive experiments are conducted to systematically evaluate the proposed model from 4 aspects: (1) Sequence and structure modeling; (2) Antigen-binding CDR-H3 generation; (3) Antibody optimization tasks; (4) Ablation analysis of key model components.

### 4.2. Data Preparation

#### SAbDab

We systematically evaluate the Sequence and structure modeling performance of RA-GHN on the Structural Antibody Database (SAbDab)[3]. After excluding structures with missing light chains or antigens, the dataset comprises a total of 3,127 antigen–antibody complexes. To reduce data redundancy and mitigate bias arising from high sequence similarity, CDR sequences are first clustered using MMseqs2[26]. Antibodies with CDR sequence identity greater than 40%, measured by the sequence identity metric[27], are assigned to the same cluster[23]. Following preprocessing, 765, 1,093, and 1,659 clusters are obtained for CDR-H1, CDR-H2, and CDR-H3, respectively, which are then split into training, validation, and test sets using an 8:1:1 ratio.

#### RAbD

To evaluate the Antigen-binding CDR-H3 generation performance of RA-GHN on antigen-related CDR-H3 design, we use the RAbD dataset[10] as an independent test benchmark, which contains 60 representative antibody– antigen complexes. Training data were curated from SAbDab with a strict data leakage prevention protocol, in which all antibodies whose CDR-H3 clusters overlapped with the RAbD test set were excluded. The remaining data were randomly split into training and validation sets at a 9:1 ratio.

#### SKEMPI V2.0

One of the central goals of antibody design is to simultaneously enhance binding affinity and binding specificity. To systematically evaluate the performance of RA-GHN in antibody optimization, we first pre-train the antibody generation model *F*_Θ_ on the SAbDab dataset, with the training and validation sets split at a ratio of 9:1. Subsequently, the pre-trained model 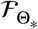 is further fine-tuned on the SKEMPI V2.0 dataset[28], which contains 53 antigen–antibody complexes, to adapt it to the antibody optimization task.

### 4.3. Implementation detail

The following hyperparameters are determined using the AutoML toolkit NNI, with the corresponding search spaces defined as follows: the Adam optimizer is adopted with a learning rate *lr* = {0.0003, 0.001}, batch size *B* = {4, 8, 16}, and training epochs *E*_train_ = 20; the embedding dimension for *E*_type_(*v*_*i*_) is set to *F*_*e*_ = 32; the hidden dimension is selected from *F* = {128, 256}, and the number of network layers is *L* = {3, 4}.

For fair comparison, we select the model checkpoint with the lowest validation loss for evaluation during testing. Moreover, all baseline methods and our approach are implemented using the standard PyTorch 1.8.0 framework. Experiments are conducted on a high-performance computing server equipped with an AMD EPYC 7742 processor (64 cores and 128 threads, base frequency 1.5 GHz and maximum boost frequency 2.25 GHz) and four NVIDIA A100 GPUs with 80 GB memory each.

### 4.4. Baselines

We compare the proposed RA-GHN with a range of representative state-of-the-art methods, including LSTM[14], RefineGNN[19], HERN [20], DiffAb[2], AbDiffuser[22], MEAN[23], dyMEAN[24] RAAD[25], and ADesigner[29]. Since LSTM, RefineGNN, and HERN model only partial contexts of the antigen–antibody complex (with LSTM and RefineGNN using only the antibody heavy chain and HERN using only the antigen), we additionally compare three extended variants—C-LSTM, C-RefineGNN, and C-HERN—by providing the complete antigen–antibody complex as input, ensuring a fair and consistent comparison across methods.

## 5. Results and Discussion

### 5.1. Sequence and Structure Modeling

We adopt Amino Acid Recovery (AAR) and Root Mean Squared Deviation (RMSD) as evaluation metrics for sequence generation and structure prediction, respectively, and report the overall results based on 10-fold cross-validation in Table 1.

**Table 1.** Average 10-fold CV performance on CDR sequence and structure modeling from SAbDab; bold and underline indicate the best and second-best values, respectively.

| Model | CDR-H1 |  | CDR-H2 |  | CDR-H3 |  |
| --- | --- | --- | --- | --- | --- | --- |
|  | AAR (↑) | RMSD (↓) | AAR (↑) | RMSD (↓) | AAR (↑) | RMSD (↓) |
| LSTM | 49.98 | - | 28.50 | - | 15.69 | - |
| C-LSTM | 40.93 | - | 29.24 | - | 15.48 | - |
| RefineGNN | 39.40 | 3.22 | 37.06 | 3.64 | 21.13 | 6.00 |
| C-RefineGNN | 33.19 | 3.25 | 33.53 | 3.69 | 18.88 | 6.22 |
| HERN | 41.71 | 1.78 | 35.46 | 2.21 | 24.75 | 3.89 |
| C-HERN | 47.36 | 1.45 | 41.45 | 1.20 | 28.47 | 2.64 |
| DiffAb | 61.34 | 1.02 | 37.66 | 1.20 | 25.79 | 3.02 |
| AbDiffuser | 62.76 | 1.08 | 41.10 | 1.16 | 29.58 | 2.83 |
| MEAN | 58.29 | 0.98 | 47.15 | 0.95 | 36.38 | 2.21 |
| dyMEAN | 63.52 | 0.84 | 55.41 | 0.82 | 37.19 | 2.09 |
| ADesigner | 64.34 | 0.82 | 55.52 | 0.79 | 37.37 | 1.97 |
| RADD | **67.64** | <u>0.77</u> | **58.87** | <u>0.72</u> | <u>37.71</u> | <u>1.91</u> |
| RA-GHN (ours) | <u>66.22</u> | **0.65** | <u>58.58</u> | **0.56** | **38.15** | **1.78** |
| $\Delta_{\text{dyMEAN}}$ | +4.25% | -22.62% | +5.72% | -31.71% | +2.58% | -14.83% |
| $\Delta_{\text{ADesigner}}$ | +2.92% | -20.73% | +5.51% | -29.11% | +2.09% | -9.64% |
| $\Delta_{\text{RADD}}$ | -2.10% | -15.58% | -0.49% | -22.22% | +1.17% | -6.81% |

As detailed in Table 1, RA-GHN achieves a superior trade-off between sequence recovery and structural accuracy, demonstrating clear advantages across biologically critical metrics. Specifically, the model attains the lowest RMSD across all CDR loops; compared with the strongest baseline, RADD, it reduces structural error by 15.58%, 22.22%, and 6.81% on CDR-H1, CDR-H2, and CDR-H3, respectively. From a biophysical perspective, this pronounced reduction in RMSD implies that the predicted structures reside closer to the global energy minimum, effectively mitigating high-energy steric clashes associated with misfolded states. Such geometric precision is strongly correlated with antibody developability, significantly reducing the risk of aggregation and instability during downstream experimental characterization.

Crucially, RA-GHN validates its effectiveness on the CDR-H3 loop, the most hypervariable region and the primary determinant of antigen-binding specificity. While canonical CDR-H1 and CDR-H2 loops are structurally conserved and amenable to statistical memorization, CDR-H3 requires modeling complex, long-range dependencies. RA-GHN achieves both the highest AAR (38.15%) and the lowest RMSD (1.78 Å) on this challenging region, confirming that the Equivariant Hypergraph Encoder effectively captures multi-body interactions beyond simple distribution fitting. Although the model exhibits a marginal decrease in AAR on the conserved CDR-H1 and CDR-H2 loops (∼2%) compared to baselines, this is outweighed by a substantial reduction in structural error (∼20%). This result highlights a strategic prioritization of functional feasibility: RA-GHN effectively bridges the gap between sequence and structure, ensuring that generated antibodies are not merely native-like in sequence but are physically realizable and structurally stable.

### 5.2. Antigen-binding CDR-H3 Generation

Modeling the hypervariable CDR-H3 region presents a fundamental dichotomy: metrics emphasizing amino acid recovery (AAR) bias models toward conservative memorization of native sequences, whereas metrics relying solely on RMSD are prone to “folding hallucination”—a failure mode in which generated sequences yield deceptively low-RMSD structures despite lacking the biophysical properties required for stability. To resolve this issue, we propose the Structure–Sequence Decoupling Index (SSDI), a composite metric designed to evaluate a model’s ability to explore novel sequence space without compromising structural physical plausibility.

Inspired by the F-measure, SSDI employs a harmonic mean to nonlinearly couple structural reliability (*P*_struct_) and sequence novelty (*P*_novel_). Unlike an arithmetic mean, the harmonic mean introduces an implicit soft-gating mechanism that requires both criteria to be satisfied simultaneously. A severe deficiency in either dimension (e.g., *P*_struct_ → 0 due to misfolding) creates a bottleneck that sharply penalizes the overall score. Formally, SSDI is defined as:

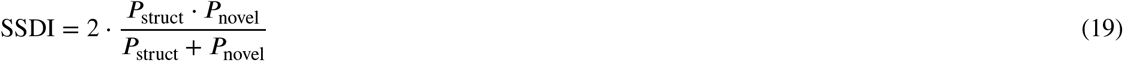

Structural reliability (*P*_struct_) maps the unbounded root-mean-square deviation (RMSD) into a probabilistic space via a sigmoid transformation:

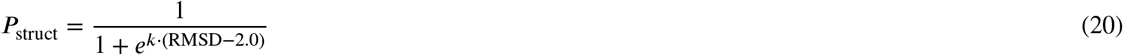

Here, the function is explicitly centered at 2.0 Å. This threshold represents a widely accepted “gold standard” in structural biology for high-accuracy structure prediction[30, 31, 32]. It delineates native-like conformations where chemical details and side-chain orientations become highly reliable—a crucial requirement during preprocessing pipelines, such as when extracting highly flexible CDR tokens for a structural antibody database. Furthermore, the parameter *k* = 3 controls the steepness of the transition. This specific value is mathematically calibrated to balance rigorous filtering with optimization stability; it establishes a soft boundary that guarantees smooth differentiability for gradient-based optimization, ensuring scores decay precipitously for physically implausible designs (RMSD > 2.0 Å) while rewarding high-precision conformations.

Sequence novelty (*P*_novel_) quantifies divergence from the native sequence. To ensure the metric captures meaningful design variations rather than statistical artifacts, it is normalized against a random baseline:

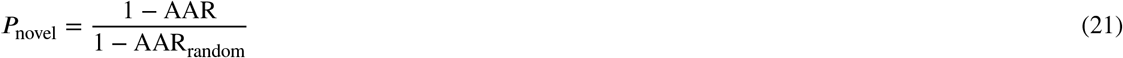

where AAR_random_ ≈ 0.05. This value reflects the statistical probability (1/20) of randomly guessing the correct standard amino acid at any given position. By subtracting this baseline, the formulation effectively eliminates the background noise of chance mutations. Under this formulation, *P*_novel_ → 1 indicates maximal, non-random sequence divergence, whereas *P*_novel_ → 0 corresponds to a near-exact reproduction of the native sequence. Crucially, within the harmonic-mean framework, this high novelty is rewarded only when accompanied by high structural reliability, thereby effectively filtering out random, physically implausible designs.

As detailed in Table 2, RA-GHN demonstrates a decisive advantage on the comprehensive SSDI metric, attaining a score of 0.7421—surpassing the second-best model, RAAD, by 4.46% and dyMEAN by 8.40%. While earlier methods (e.g., RosettaAD and RefineGNN) yield near-zero scores due to a strict soft-gating mechanism that penalizes imbalances between structural recovery and sequence novelty, RA-GHN successfully decouples sequence design from rigid natural imitation. This improvement validates the effectiveness of our framework in managing the critical trade-off between structural stability and sequence exploration.

**Table 2.** Performance comparison of antigen-binding CDR-H3 sequence and structure design on the RAbD dataset.

| Model | SSDI (↑) | TM-score (↑) |
| --- | --- | --- |
| RosettaAD | 0.0001 | 0.9435 |
| RefineGNN | 0 | 0.8308 |
| C-RefineGNN | 0 | 0.8317 |
| HERN | 0.0641 | 0.9684 |
| C-HERN | 0.1521 | 0.9734 |
| MEAN | 0.6519 | 0.9812 |
| dyMEAN | 0.6846 | 0.9825 |
| ADesigner | 0.6974 | <u>0.9850</u> |
| RAAD | <u>0.7104</u> | **0.9874** |
| RA-GHN (Ours) | **0.7421** | 0.9840 |
| $\Delta_{\text{dyMEAN}}$ | +8.40% | +0.15% |
| $\Delta_{\text{ADesigner}}$ | +6.41% | -0.10% |
| $\Delta_{\text{RAAD}}$ | +4.46% | -0.34% |

Although RA-GHN exhibits a marginal decrease in TM-score (∼0.34%) compared to the top-performing baseline, all generated structures maintain high biophysical plausibility, with TM-scores exceeding 0.98. This outcome reflects a deliberate design strategy: by relaxing strict local mimicry, the model gains the flexibility to discover novel sequence motifs that reliably fold into stable backbone conformations. Visual validation on representative cases (Figure 3) corroborates these quantitative results, confirming that RA-GHN prevents “folding hallucination” and faithfully reconstructs backbone orientations even for complex targets.

**Figure 3.**
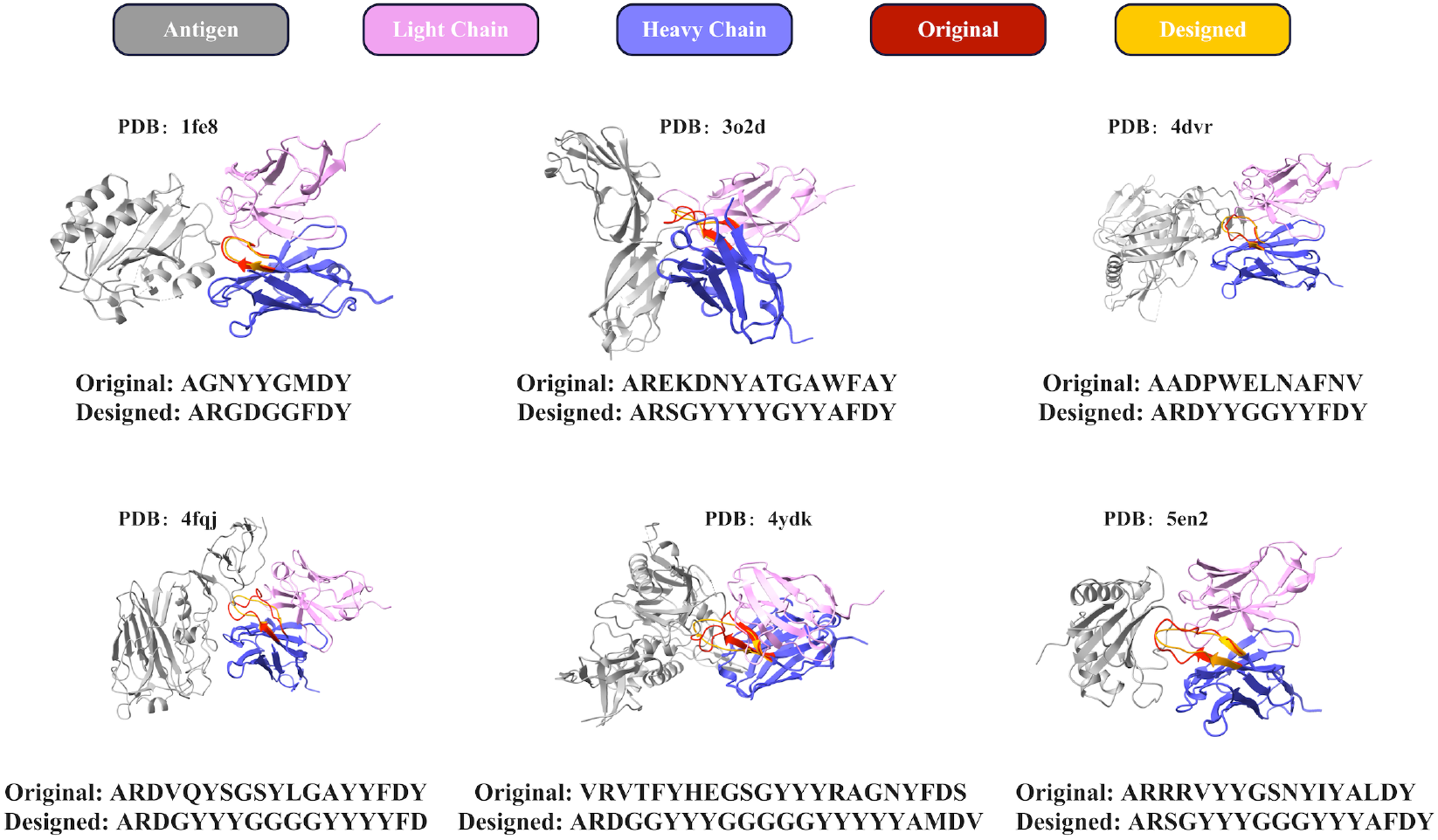
Case Study. The structures and sequences of the antigen-binding CDR-H3s designed by RA-GHN for 6 complexes.

### 5.3. Antibody Optimization

Table 3 reports the average changes in binding free energy of the optimized antibodies toward the target antigen and non-target antigens, denoted as ΔΔ*G*_pos_ and ΔΔ*G*_neg_, respectively, as well as their difference, which defines the specificity metric (SP-score).

**Table 3.** Average ΔΔ*G* (kcal/mol) after antibody optimization.

| Model | $\Delta\Delta G_{\text{pos}}$ (↓) | $\Delta\Delta G_{\text{neg}}$ (↑) | SP-score (↑) |
| --- | --- | --- | --- |
| Random | +1.489 | **+1.518** | 0.029 |
| LSTM | -2.183 | -0.496 | 1.687 |
| C-LSTM | -2.534 | -0.784 | 1.750 |
| RefineGNN | -4.684 | -1.191 | 3.493 |
| C-RefineGNN | -5.179 | -1.392 | 3.787 |
| MEAN | -6.104 | -1.605 | 4.499 |
| dyMEAN | -8.215 | -1.543 | 6.672 |
| ADesigner | -10.431 | -1.624 | 8.807 |
| RAAD | <u>-11.787</u> | <u>-0.238</u> | **11.549** |
| RA-GHN | **-12.145** | -1.213 | <u>10.932</u> |

As presented in Table 3, RA-GHN achieves state-of-the-art performance in affinity maturation, attaining the lowest average ΔΔ*G*_pos_ of −12.145 kcal/mol. This result represents a substantial improvement over both sequence-based methods (e.g., LSTM) and early graph-based models (e.g., RefineGNN). Compared with the strong competitor RAAD, RA-GHN demonstrates superior capability in minimizing binding free energy (−**12.145** vs. −11.787 kcal/mol). Although RAAD achieves a marginally higher SP-score due to its conservative behavior on negative controls, RA-GHN strategically prioritizes the primary objective of maximizing antigen-binding affinity, thereby delivering antibodies with tighter and more specific interactions.

To provide an intuitive assessment of RA-GHN’s efficacy in antibody affinity optimization, we conducted an in-depth analysis on six representative cases selected from the SKEMPI V2.0 dataset. As shown in Fig. 4, we compared the native CDR-H3 loops (red) with the RA-GHN-designed variants (Orange), focusing on structural deviations, sequence composition, and changes in ΔΔ*G*.

**Figure 4.**
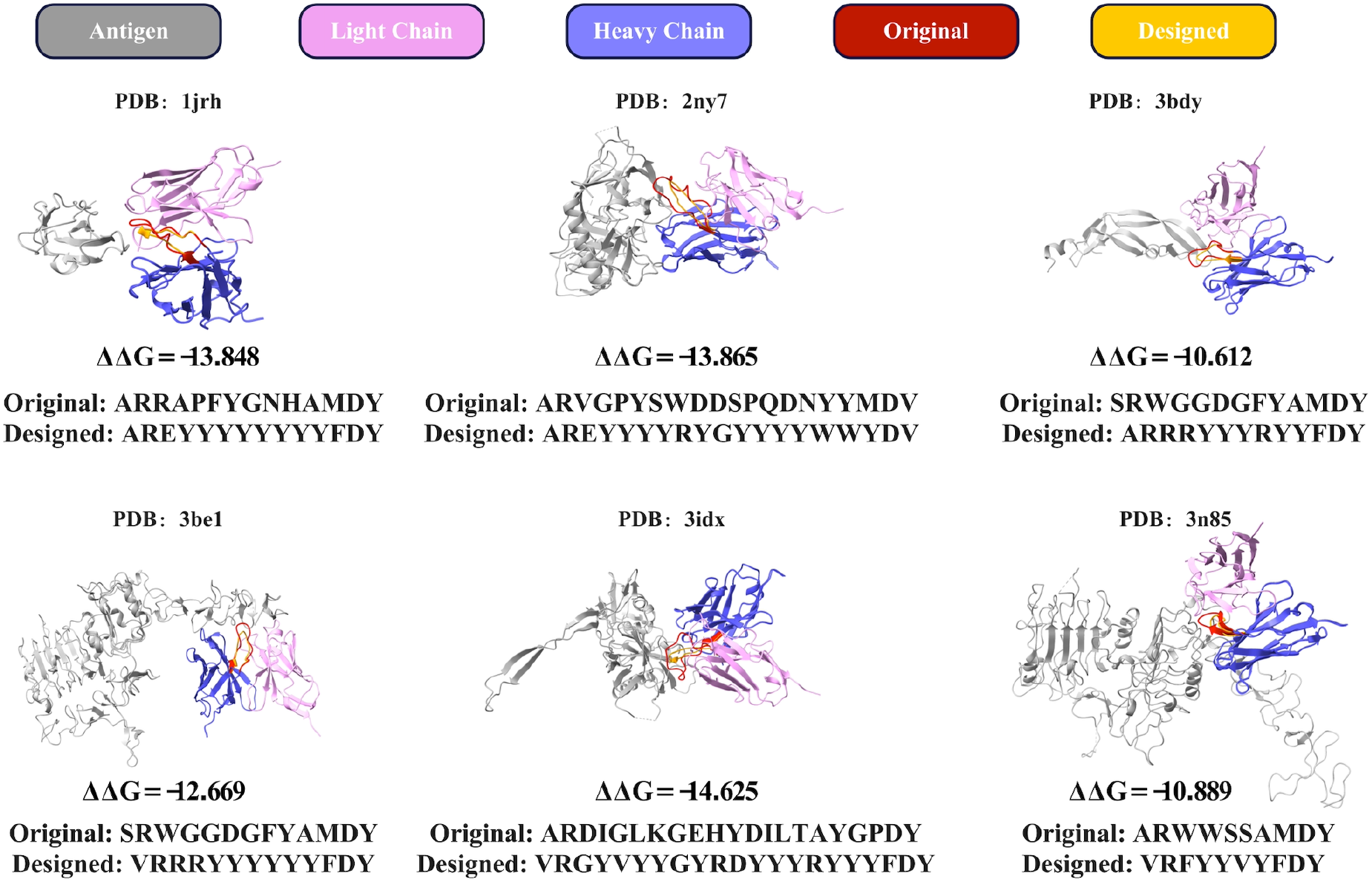
Visualization of RA-GHN-based antibody optimization on SKEMPI V2.0 test cases.

The observed affinity gains are driven by specific physicochemical and structural refinements. At the sequence level, RA-GHN identifies an enrichment of tyrosine and arginine residues within CDR-H3 loops, optimizing interface stabilization through *π*–*π* stacking and cation–*π* interactions. Structurally, while the design preserves the global topological framework, it introduces adaptive local refinements that enhance geometric complementarity to the epitope. Notably, in cases such as the 2ny7 complex, the model proposes a distinct physicochemical profile while maintaining the canonical backbone conformation. This observation confirms that RA-GHN actively navigates the fitness landscape to discover novel, high-affinity solutions rather than merely memorizing evolutionary history.

### 5.4. Ablation Study

#### Ablation study on hyperedge modeling strategies

We systematically conduct an ablation study on the sequence and structure modeling task to evaluate the impact of incorporating hyperstructures and to analyze the efficacy of distinct hyperedge combinations. The experimental settings span a comprehensive spectrum, ranging from single-view modeling to multi-view fusion.

As illustrated in Fig. 5, the ablation study elucidates the critical interplay between functional modeling and geometric constraints. The results indicate that explicit epitope modeling is the primary driver of sequence recovery: the Epitope-only setting achieves a remarkable AAR of 38.10%, confirming that paratope–epitope interactions provide the decisive signal for decoding binding specificity. Conversely, relying solely on geometric constraints (Spatial+Hub) leads to over-regularization; although this configuration minimizes RMSD (1.778 Å), it incurs a noticeable drop in AAR (37.12%). This observation suggests that purely geometric optimization tends to “over-smooth” sequence distributions, thereby discarding functionally essential residues that may be structurally non-ideal.

**Figure 5.**
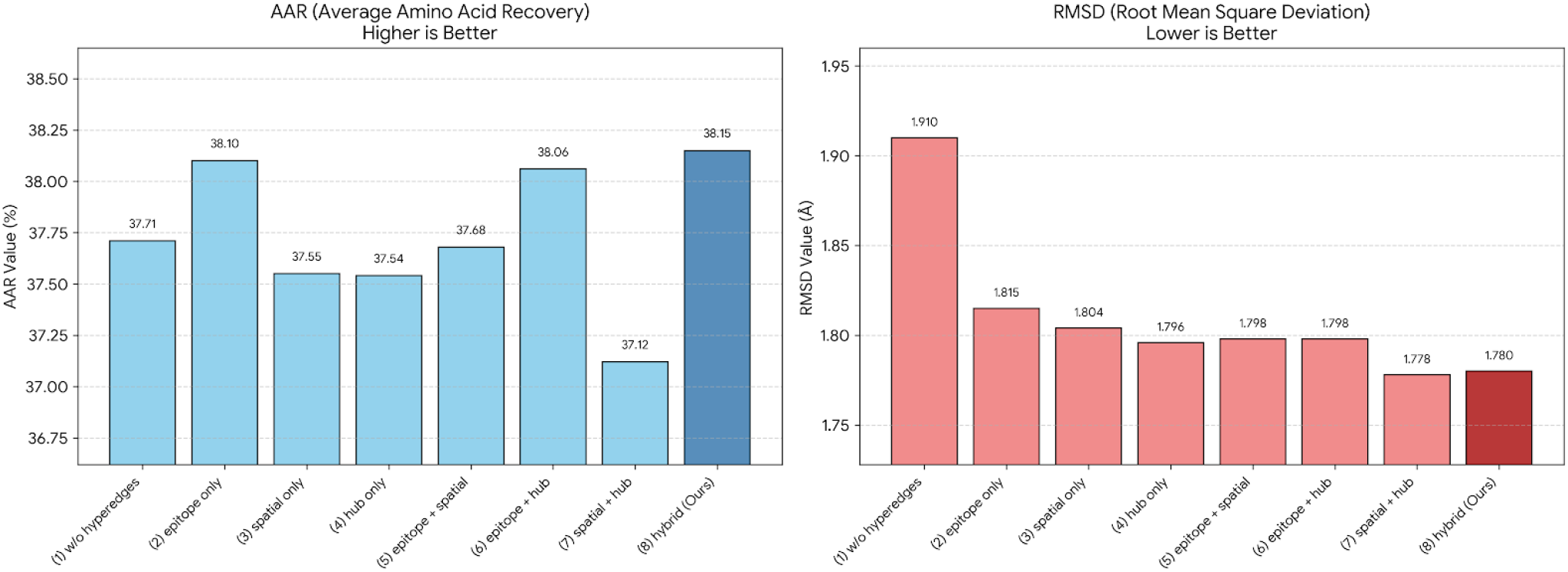
Ablation study on hyperedge modeling strategies. The bar charts compare sequence recovery (AAR, left) and structural accuracy (RMSD, right) across different configurations.

The Hybrid (RA-GHN) framework effectively resolves this tension. By fusing the semantic guidance of the epitope view with the physical precision of the spatial-hub views, it achieves the best overall performance (AAR: 38.15%, RMSD: 1.780 Å). This result demonstrates that our multi-view hypergraph does not merely trade off between competing objectives, but instead synergistically optimizes both biological function and physical structural integrity.

#### Sensitivity Analysis of the Balancing Hyperparameter α

We conducted a sensitivity analysis on α, the weight of the coordinate loss, to determine the optimal balance between sequence generation and structure prediction (Figure 6). Initially, increasing α from 0.1 to 0.2 yields a synergistic improvement, culminating in a peak AAR of 38.15% and a competitive RMSD of 1.778 Å at α = 0.2. This result suggests that moderate structural supervision acts as a beneficial regularizer, pruning biologically implausible conformations and thereby facilitating accurate sequence decoding.

**Figure 6.**
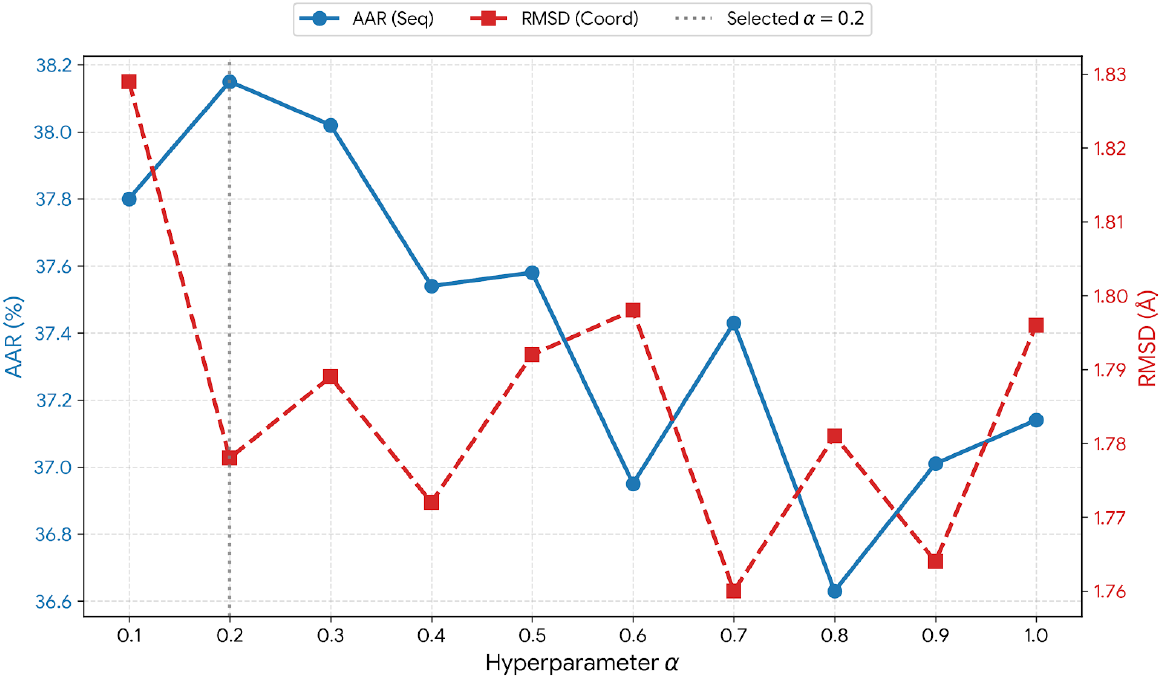
Sensitivity analysis revealing the sequence-structure trade-off.

However, a divergence emerges beyond α = 0.3. Although α = 0.7 achieves the lowest RMSD (1.760 Å), it incurs a significant drop in AAR (37.43%). This decline indicates structural over-regularization: excessive coordinate supervision drives the model toward rigid, “safe” mean-field conformations, suppressing the subtle, energetically driven sequence variations required for specific antigen binding. Consequently, we select α = 0.2 as the optimal configuration, prioritizing binding specificity (AAR) while maintaining robust structural precision.

## 6. Conclusion

In this work, we presented RA-GHN, a novel Relation-Aware Graph–Hypergraph Network designed to systematically address the challenges in antibody sequence–structure co-design. By fusing pairwise relation graphs with biophysical hypergraphs, RA-GHN explicitly models complex multi-body interactions and global geometric constraints at the antibody–antigen interface, effectively overcoming the limitations of conventional pairwise methods. The architecture employs a dual-pathway mechanism that coordinates the optimization of residue representations and three-dimensional coordinates while strictly ensuring *SE*(3)-equivariance. Furthermore, to mitigate the “folding hallucination” issue prevalent in generative antibody design, we introduce the Structure–Sequence Decoupling Index (SSDI). This metric balances physical reliability with sequence novelty, providing a rigorous and discriminative criterion for model assessment. Extensive experiments on SAbDab, RAbD, and SKEMPI demonstrate that RA-GHN significantly outperforms state-of-the-art baselines in CDR modeling, antigen-binding structure generation, and affinity optimization, confirming that incorporating high-order geometric constraints yields antibodies that are both physically feasible and functionally potent.

Despite these advancements, several avenues for further exploration remain. Future work will extend the current static modeling paradigm to account for conformational dynamics and induced-fit mechanisms. In addition, we plan to generalize the proposed graph–hypergraph architecture to broader protein design tasks, such as nanobody engineering, and to incorporate multi-objective constraints (e.g., solubility and immunogenicity). Ultimately, coupling this computational framework with high-throughput wet-lab screening will be essential for realizing a closed-loop design cycle that translates theoretical predictions into clinically viable therapeutics.

## CRediT authorship contribution statement

**Jinzhu Wei:** Writing – original draft,Writing – review & editing,Visualization, Validation, Resources, Methodology, Formal analysis. **Shuijun Zhang:** Writing – review & editing,Formal analysis. **Hongliang Duan:** Writing – review & editing,Validation,Investigation. **Leyi Wei:** Writing – review & editing, Investigation. **Jingjing Guo:** Writing – original draft, Writing – review & editing, Project administration, Funding acquisition, Formal analysis, Conceptualization.

## Notes

⋆ This work is supported by the Science and Technology Development Fund of Macau (0177/2023/RIA3).

### Competing Interest Statement

The authors have declared no competing interest.

## References

[1] D. Akpinaroglu, J. A. Ruffolo, S. P. Mahajan, J. J. Gray, Simultaneous prediction of antibody backbone and side-chain conformations with deep learning, PLOS one 17 (6) (2022) e0258173.

[2] S. Luo, Y. Su, X. Peng, S. Wang, J. Peng, J. Ma, Antigen-specific antibody design and optimization with diffusion-based generative models for protein structures, in: Advances in Neural Information Processing Systems (NeurIPS), Vol. 35, 2022, pp. 9754–9767.

[3] R. Akbar, P. A. Robert, C. R. Weber, M. Widrich, R. Frank, M. Pavlović, L. Scheffer, M. Chernigovskaya, I. Snapkov, A. Slabodkin, et al., In silico proof of principle of machine learning-based antibody design at unconstrained scale, in: MAbs, Vol. 14, Taylor & Francis, 2022, p. 2031482.

[4] J. Dunbar, K. Krawczyk, J. Leem, T. Baker, A. Fuchs, G. Georges, J. Shi, C. M. Deane, Sabdab: the structural antibody database, Nucleic acids research 42 (D1) (2014) D1140–D1146.

[5] K. Murphy, C. Weaver, Janeway’s immunobiology, Garland science, 2016.

[6] D. T. Chan, M. A. Groves, Affinity maturation: highlights in the application of in vitro strategies for the directed evolution of antibodies, Emerging Topics in Life Sciences 5 (5) (2021) 601–608.

[7] M. I. Raybould, C. Marks, K. Krawczyk, B. Taddese, J. Nowak, A. P. Lewis, A. Bujotzek, J. Shi, C. M. Deane, Five computational developability guidelines for therapeutic antibody profiling, Proceedings of the National Academy of Sciences 116 (10) (2019) 4025–4030.

[8] R. Pantazes, C. D. Maranas, Optcdr: a general computational method for the design of antibody complementarity determining regions for targeted epitope binding, Protein Engineering, Design & Selection 23 (11) (2010) 849–858.

[9] G. D. Lapidoth, D. Baran, G. M. Pszolla, C. Norn, A. Alon, M. D. Tyka, S. J. Fleishman, Abdesign: A n algorithm for combinatorial backbone design guided by natural conformations and sequences, Proteins: Structure, Function, and Bioinformatics 83 (8) (2015) 1385–1406.

[10] J. Adolf-Bryfogle, O. Kalyuzhniy, M. Kubitz, B. D. Weitzner, X. Hu, Y. Adachi, W. R. Schief, R. L. Dunbrack Jr, Rosettaantibodydesign (rabd): A general framework for computational antibody design, PLoS computational biology 14 (4) (2018) e1006112.

[11] S. Warszawski, A. Borenstein Katz, R. Lipsh, L. Khmelnitsky, G. Ben Nissan, G. Javitt, O. Dym, T. Unger, O. Knop, S. Albeck, et al., Optimizing antibody affinity and stability by the automated design of the variable light-heavy chain interfaces, PLoS computational biology 15 (8) (2019) e1007207.

[12] M. M. Gomari, P. Tarighi, E. Choupani, S. Abkhiz, M. Mohamadzadeh, N. Rostami, E. Sadroddiny, S. Baammi, V. N. Uversky, N. V. Dokholyan, Structural evolution of delta lineage of sars-cov-2, International journal of biological macromolecules 226 (2023) 1116–1140.

[13] J.-E. Shin, A. J. Riesselman, A. W. Kollasch, C. McMahon, E. Simon, C. Sander, A. Manglik, A. C. Kruse, D. S. Marks, Protein design and variant prediction using autoregressive generative models, Nature communications 12 (1) (2021) 2403.

[14] K. Saka, T. Kakuzaki, S. Metsugi, D. Kashiwagi, K. Yoshida, M. Wada, H. Tsunoda, R. Teramoto, Antibody design using lstm based deep generative model from phage display library for affinity maturation, Scientific reports 11 (1) (2021) 5852.

[15] B. Jing, G. Corso, J. Chang, R. Barzilay, T. Jaakkola, Torsional diffusion for molecular conformer generation, in: Advances in neural information processing systems (NeurIPS), Vol. 35, 2022, pp. 24240–24253.

[16] B. Lai, M. McPartlon, J. Xu, End-to-end deep structure generative model for protein design, bioRxiv (2022). doi:10.1101/2022.07.09.499440.

[17] R. Akbar, H. Bashour, P. Rawat, P. A. Robert, E. Smorodina, T.-S. Cotet, K. Flem-Karlsen, R. Frank, B. B. Mehta, M. H. Vu, et al., Progress and challenges for the machine learning-based design of fit-for-purpose monoclonal antibodies, in: MAbs, Vol. 14, Taylor & Francis, 2022, p. 2008790.

[18] E. C. Alley, G. Khimulya, S. Biswas, M. AlQuraishi, G. M. Church, Unified rational protein engineering with sequence-based deep representation learning, Nature methods 16 (12) (2019) 1315–1322.

[19] W. Jin, J. Wohlwend, R. Barzilay, T. S. Jaakkola, Iterative refinement graph neural network for antibody sequence-structure co-design, in: International Conference on Learning Representations (ICLR), 2022, pp. 1–14.

[20] W. Jin, R. Barzilay, T. Jaakkola, Antibody-antigen docking and design via hierarchical structure refinement, in: International Conference on Machine Learning (ICML), PMLR, 2022, pp. 10217–10227.

[21] E. Hoogeboom, D. Nielsen, P. Jaini, P. Forré, M. Welling, Argmax flows and multinomial diffusion: Learning categorical distributions, in: Advances in neural information processing systems (NeurIPS), Vol. 34, 2021, pp. 12454–12465.

[22] K. Martinkus, J. Ludwiczak, W.-C. Liang, J. Lafrance-Vanasse, I. Hotzel, A. Rajpal, Y. Wu, K. Cho, R. Bonneau, V. Gligorijevic, et al., Abdiffuser: full-atom generation of in-vitro functioning antibodies, in: Advances in Neural Information Processing Systems (NeurIPS), Vol. 36, 2023, pp. 40729–40759.

[23] X. Kong, W. Huang, Y. Liu, Conditional antibody design as 3d equivariant graph translation, in: The Eleventh International Conference on Learning Representations (ICLR), 2022, pp. 1–21.

[24] X. Kong, W. Huang, Y. Liu, End-to-end full-atom antibody design, in: International Conference on Machine Learning (ICML), PMLR, 2023, pp. 17409–17429.

[25] L. Wu, H. Lin, Y. Huang, Z. Gao, C. Tan, Y. Liu, T. Wu, S. Z. Li, Relation-aware equivariant graph networks for epitope-unknown antibody design and specificity optimization, in: Proceedings of the AAAI Conference on Artificial Intelligence (AAAI), Vol. 39, 2025, pp. 895–904.

[26] M. Steinegger, J. Söding, Mmseqs2 enables sensitive protein sequence searching for the analysis of massive data sets, Nature biotechnology 35 (11) (2017) 1026–1028.

[27] S. Henikoff, J. G. Henikoff, Amino acid substitution matrices from protein blocks., Proceedings of the national academy of sciences 89 (22) (1992) 10915–10919.

[28] J. Jankauskaitė, B. Jiménez-García, J. Dapkūnas, J. Fernández-Recio, I. H. Moal, Skempi 2.0: an updated benchmark of changes in protein– protein binding energy, kinetics and thermodynamics upon mutation, Bioinformatics 35 (3) (2019) 462–469.

[29] C. Tan, Z. Gao, L. Wu, J. Xia, J. Zheng, X. Yang, Y. Liu, B. Hu, S. Z. Li, Cross-gate mlp with protein complex invariant embedding is a one-shot antibody designer, in: Proceedings of the AAAI Conference on Artificial Intelligence (AAAI), Vol. 38, 2024, pp. 15222–15230.

[30] J. Jumper, R. Evans, A. Pritzel, T. Green, M. Figurnov, O. Ronneberger, K. Tunyasuvunakool, R. Bates, A. Žídek, A. Potapenko, et al., Highly accurate protein structure prediction with alphafold, Nature 596 (7873) (2021) 583–589.

[31] J. C. Almagro, A. Teplyakov, J. Luo, R. W. Sweet, S. Kodangattil, F. Hernandez-Guzman, G. L. Gilliland, Second antibody modeling assessment (ama-ii) (2014).

[32] J. A. Ruffolo, L.-S. Chu, S. P. Mahajan, J. J. Gray, Fast, accurate antibody structure prediction from deep learning on massive set of natural antibodies, Nature communications 14 (1) (2023) 2389.

